# HAETAE: A highly accurate and efficient epigenome transformer for tissue-specific histone modification prediction

**DOI:** 10.64898/2026.03.09.708388

**Authors:** Seung-Jin Park, Soo-Hyun Im, Kim Jae-Yoon, Seon-Young Kim

## Abstract

While genomic models trained on four bases often fail to capture cell-type specificity, we introduce HAETAE, which integrates 5-methylcytosine from long-read sequencing into a 5-base framework. By explicitly modeling epigenetic context, HAETAE achieves state-of-the-art accuracy (>0.95) with orders of magnitude fewer parameters, challenging the prevailing scaling-law paradigm. Furthermore, HAETAE deciphers tissue-specific regulatory logic, as demonstrated by revealing the distinct, context-dependent functional impact of the TERT promoter mutation across diverse tissues.

## Main

Although cells share an identical genome, their distinct identities are defined by how these genomic instructions are differentially regulated^1^. Recent breakthroughs in deep learning have introduced powerful sequence-to-function architectures—such as Enformer^2^ and AlphaGenome^3^, alongside genomic language models like Nucleotide Transformer (NT)^4^, DNABERT^5^, and HyenaDNA^6^ —which have demonstrated remarkable success in predicting gene expression and assessing variant effects. However, by relying on a static four-nucleotide code, these models remain insufficient for capturing the tissue-specific regulatory mechanisms^7–9^. Addressing this limitation requires integrating explicit regulatory signals; here, we identify 5-methylcytosine (5mC) as a critical determinant for capturing these tissue-specific epigenetic landscapes. HAETAE was developed by incorporating methylation status directly into the model vocabulary (Fig. 1a). By leveraging long-read sequencing data across three tissues (Supplementary Table. 1), HAETAE encodes methylation status as a distinct fifth token (‘M’). This 5-base architecture enables the efficient learning of tissue-specific regulatory syntax with only 0.2 million parameters (Supplementary Fig. 1), thereby facilitating more precise modeling of regulatory mechanisms, such as prediction of histone ChIP-seq peaks.

**Fig. 1.**
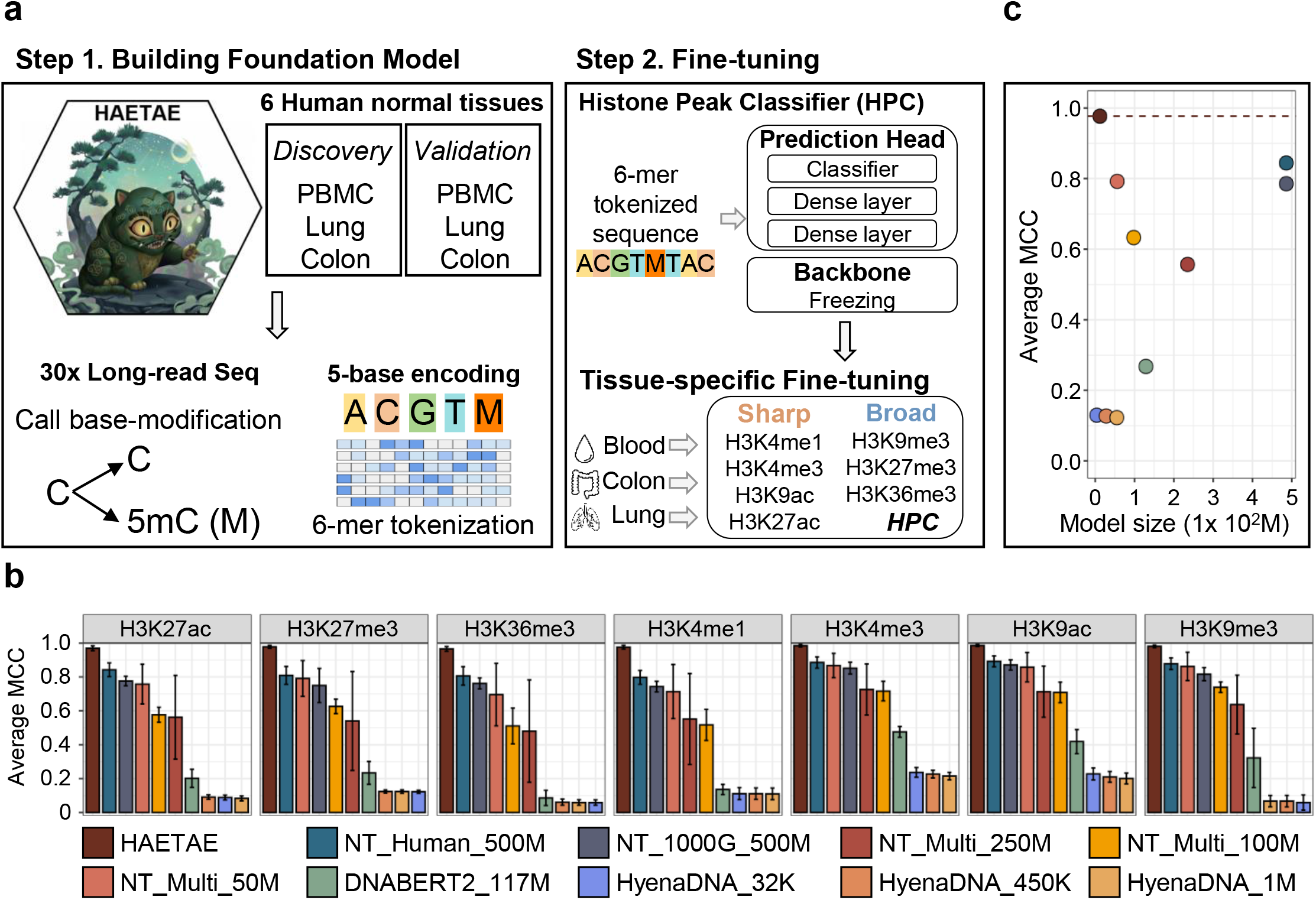
HAETAE framework and efficiency. a, Schematic of the 5-base modeling approach. b, Performance benchmarking against foundation models across seven histone marks. c, Accuracy relative to model parameters.

Benchmarking the prediction of peaks for seven histone modifications across three tissues from ENCODE data (Supplementary Fig. 2) revealed that HAETAE significantly outperforms several 4-base models, including NT, DNABERT2, and HyenaDNA, achieving superior predictive power in all conditions (Fig. 1b, Supplementary Fig. 3, Supplementary Table 2). This superior performance was consistent across multiple metrics, including MCC, AUROC, and F1 score (Supplementary Table 3).

Notably, HAETAE achieved an accuracy of >0.95 with a highly compact architecture, demonstrating exceptional parameter efficiency (Fig. 1c). This result underscores a key principle of data-centric AI: the informational density of the training data is often more critical than sample size. By integrating high-fidelity methylation signals (5mC) from deep-coverage (~30x) long-reads, HAETAE captures tissue-specific regulatory logic more effectively than models trained on massive but static 4-base datasets. This suggests that explicit epigenetic priors provide a decisive advantage over simply scaling sequence volume.

To validate the contribution of the methylation token, we performed ablation studies. Replacing ‘M’ with ‘C’ (M>C) reduced performance to the 0.7–0.8 range, comparable to baseline models (Fig. 2a), a trend consistent across all tested tissues and histone marks (Supplementary Fig. 4 and Supplementary Table. 4). Attention analysis indicated that the model assigns higher weights to 5mC in the core regions of peaks compared to flanking boundaries, accurately reflecting the positional importance of methylation in defining ChIP-seq peaks (Supplementary Fig. 5 and Supplementary Table. 5). This finding aligns with our input window design (peak summit ±500 bp), confirming that the model effectively learns positional importance. Conversely, a linear logistic regression baseline using simple 5-base counts yielded an average MCC of approximately 0.2 (Supplementary Table. 6). This result indicates that HAETAE does not merely rely on the density of CpG sites or methylation frequency. Instead, HAETAE learns high-order sequence context—such as biological syntax and genomic dependencies—and the complex interplay between sequence and epigenetics, distinguishing true regulatory peaks from simple genomic repeats.

**Fig. 2.**
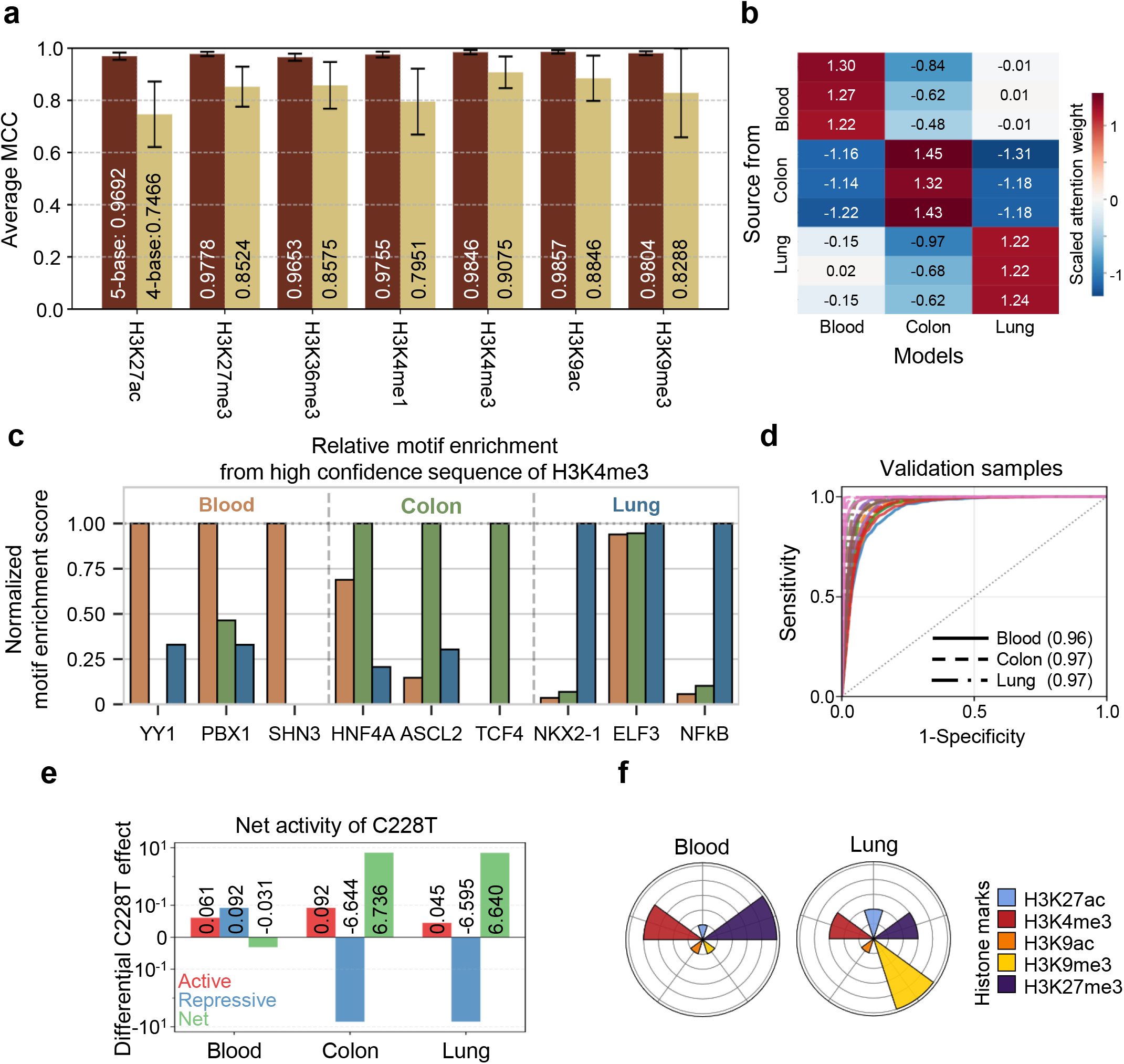
Tissue specificity and variant interpretation. a, Performance impact of M>C ablation (mean ± s.d.). b, Prediction specificity under tissue-mismatched profiles. c, Lineage-specific motif enrichment in H3K4me3 peaks. d, Generalization (AUC) on independent samples. e, Predicted TERT C228T activity across tissues. f, Differential changes from C228T.

We further assessed the model’s ability to capture tissue specificity, finding that prediction probabilities dropped significantly when sequences were tested with mismatched epigenomic profiles, such as placing a lung methylation pattern on a colon sequence (Fig. 2b). To further validate this dependency on precise epigenetic contexts, we performed robustness tests showing that while performance degrades when ‘M’ tokens are randomized, the model retains some predictive power, indicating it has learned to distinguish contextually relevant methylation from noise (Supplementary Fig. 6).

The model’s biological relevance was confirmed through motif enrichment analysis. HAETAE correctly identified nine lineage-specific transcription factor binding motifs within high-confidence predictions (Fig. 2c). In a detailed analysis of colon tissue, the model’s predicted motif activities showed high concordance with known regulatory roles (Supplementary Fig. 7). For instance, the model identified THAP11 and ZSCAN4C as the top activator and repressor, respectively, while correctly prioritizing tissue-specific markers like HNF4A and ASCL2 within the top tier of evaluated motifs. This demonstrates that HAETAE can prioritize biologically relevant regulators in a tissue-specific manner. Additionally, the model showed high generalization capability (AUC = 0.97) when tested on independent samples processed with the same pipeline (Fig. 2d).

Finally, we applied HAETAE to dissect the TERT C228T mutation. Consistent with the known landscape of TERT regulation, the model predicted a substantial activating effect specifically in solid tissues (lung and colon) compared to the minimal activity observed in blood (Fig. 2e). Beyond successfully predicting the activation of the TERT C228T mutation in solid tissues, HAETAE deconvolved the underlying chromatin syntax driving tissue specificity. Specifically, the model accurately attributed upregulation in lung tissue to H3K4me3 gain against a repressive H3K9me3 baseline, recapitulating established epigenetic landscapes^10^. This capacity to isolate context-dependent drivers demonstrates HAETAE’s potential for the precise, mechanistic interpretation of non-coding variants.

In conclusion, HAETAE demonstrates that explicit modeling of epigenetic context via a 5-base framework challenges the prevailing reliance on massive parameter scaling. This context-aware approach not only bridges the genotype-phenotype gap but also offers unprecedented experimental efficiency: a single long-read WGS run can now decipher comprehensive regulatory layers, effectively serving as a scalable alternative to parallel ChIP-seq profiling. As long-read sequencing becomes standard, HAETAE establishes a scalable, data-centric blueprint for next-generation genomic foundation models.

## Supplementary Figure legends

Supplementary Fig. 1. HAETAE model architecture and pre-training. Detailed schematic of the model architecture, including the embedding layer for the 5-base vocabulary, attention mechanisms, and the pre-training objective. The model comprises approximately 0.2 million learnable parameters.

Supplementary Fig. 2. Benchmarking of HAETAE against existing genomic foundation models. Performance comparison using Matthews Correlation Coefficient (MCC) across seven histone marks in three tissues (Blood, Colon, Lung). HAETAE consistently outperforms other state-of-the-art models, including Nucleotide Transformer (NT) variants, DNABERT2, and HyenaDNA. Error bars represent mean ± s.d. from three independent runs.

Supplementary Fig. 3. Robustness of performance across models. Matthews Correlation Coefficient (MCC) comparison of 10 genomic models (HAETAE, 5 NT variants, DNABERT, HyenaDNA) across 3 tissues and 7 histone marks. Data represent the mean of three independent training runs. HAETAE shows superior MCC across all conditions.

Supplementary Fig. 4. Comprehensive ablation study results. Detailed breakdown of the M-ablation test (M>C) for all 3 tissues and 7 histone modifications. While the magnitude of the drop varies, the 5-base input yields higher performance in every distinct case compared to the cytosine-only input.

Supplementary Fig. 5. Predicted motif activity landscape in colon tissue. The central plot displays the ranked net activity of transcription factor motifs, derived from the aggregation of predicted active and repressive histone marks. Detailed panels illustrate the specific histone modification profiles for the top-ranked activator (THAP11), the most repressed motif (ZSCAN4C), and key colon-specific lineage markers (HNF4A and ASCL2), demonstrating the model’s ability to prioritize biologically relevant regulators.

Supplementary Fig. 6. Robustness against methylation noise. Performance degradation analysis when ‘M’ tokens are assigned randomly versus derived from ML scores. H3K9me3 prediction shows the most gradual decline, while H3K4me3 shows the sharpest drop, indicating varying degrees of sensitivity to epigenetic context.

Supplementary Fig. 7. Motif activity analysis in colon tissue. Ranked motif net activity predictions in colon tissue. a, THAP11 is identified as the most activated motif, while ZSCAN4C is the most repressed. b, Tissue-specific markers ASCL2 and HNF4A rank 80th and 117th, respectively, among ~600 motifs, validating the model’s biological relevance for gene expression prediction.

## Supplementary Table legends

Supplementary Table 1. Sample information and quality control. Summary of the six PacBio WGS samples used in this study, including tissue of origin, sequencing depth, and basic quality control metrics.

Supplementary Table 2. Extended performance evaluation of HAETAE and benchmark models. Detailed performance metrics (including MCC, AUROC, AUPRC, F1 score, and Accuracy) comparing HAETAE against baseline genomic models across three tissues (Blood, Colon, Lung) and seven histone marks. The table includes individual scores from three independent training seeds to demonstrate model stability and reproducibility.

Supplementary Table 3. Extended performance metrics. Comparison of HAETAE against benchmark models using AUROC, AUPRC, F1 score, and Accuracy. HAETAE demonstrates superior performance across all evaluated metrics.

Supplementary Table 4. Quantitative ablation study on the impact of methylation tokenization. Performance comparison (MCC) between the standard HAETAE model (5-base input) and an ablated version where 5-methylcytosine tokens were replaced with cytosine (4-base input; M>C). The consistent performance drop observed in the ablated model across all evaluated tissues and histone marks quantifies the critical contribution of explicit epigenetic modeling to prediction accuracy.

Supplementary Table 5. Positional weight analysis of 5mC. Analysis of attention weights assigned to 5mC tokens in the core (center) versus boundary (flank) regions of 1000bp sequences. Higher weights in the core region confirm the model correctly learns the positional importance of methylation in defining peak centers.

Supplementary Table 6. Performance of linear logistic regression baseline. MCC scores for a simple logistic regression model using counts of A, C, G, T, and M. The failure of this model to exceed MCC > 0.5 indicates that HAETAE’s performance derives from learning high-order sequence context rather than simple nucleotide frequency.

## Online content

Any methods, additional references, Nature Portfolio reporting summaries, supplementary information, acknowledgements, peer review information; details of author contributions and competing interests; and statements of data and code availability are available at TBA.

## Notes

### Competing Interest Statement

The authors have declared no competing interest.

